# Phylogenetic Distribution of Prokaryotic Non-Homologous End Joining DNA Repair Systems in Bacteria and Archaea

**DOI:** 10.1101/2023.09.30.560322

**Authors:** Femila Manoj, Thomas E. Kuhlman

## Abstract

Nonhomologous end-joining (NHEJ) is a repair mechanism for double strand breaks (DSBs) of DNA. This mechanism is ubiquitously observed within the eukaryotic domain; however, its presence is not as pervasive among prokaryotes and archaea. Notably, in prokaryotes, it has been discerned that multiple distinct NHEJ pathways have evolved in contrast to the singular NHEJ pathway prevalent in eukaryotes. We performed phylogenetic analysis to gain deeper insights into the distribution of these prokaryotic NHEJ pathways. Concurrently, components of the prokaryotic NHEJ pathways were used to find if any archaea carry the genes required and may be able to carry out NHEJ. The results show that few prokaryotes carry the components required for NHEJ, but multiple pathways may be active in a single species. In the context of Archaea, the analysis revealed that a substantial number of species contain fragments or segments of prokaryotic NHEJ elements. Nevertheless, the presence of all the necessary components for the complete execution of the NHEJ pathway remains relatively rare within the archaeal domain.

## INTRODUCTION

Non-homologous end joining (NHEJ) is a crucial repair pathway that safeguards genome integrity by resolving DNA double-strand breaks (DSBs) when a suitable DNA template for homologous recombination (HR) is lacking ^1^. While the basic mechanisms of NHEJ have been explained, further investigations have uncovered numerous NHEJ-like genes in bacterial genomes, suggesting the existence of other DNA repair components ^2^. Additionally, prokaryotic NHEJ is relatively scarce in Archaea, with key NHEJ proteins, such as Ku, absent in most Archaea ^3^. While all the genes for proteins required for prokaryotic NHEJ can be found in very few Archaea, the genes are not next together in the genome and do not form a single NHEJ operon. Instead, the individual NHEJ-associated genes are found spread out around the genome ^4^.

While all eukaryotes can perform NHEJ, the identification of prokaryotes able to perform NHEJ has been recent. Prokaryotic NHEJ was first discovered in *Bacillus subtilis* 21 years ago ^5^. This involved targeted deletion of the genes responsible for LigD and Ku proteins, followed by assessing the survival rate after exposure to ionizing radiation, which induces DSBs. Prior to this discovery, it was widely believed that prokaryotes exclusively relied on HR for DNA repair. Subsequently, it was demonstrated that *Mycobacterium tuberculosis* and *Mycobacterium smegmatis* possess an operational NHEJ pathway capable of repairing plasmid DNA following transformation.^6^ This further solidified the notion that numerous prokaryotic species possess the ability to employ NHEJ for DNA repair processes. However, many prokaryotes, such as *Escherichia coli*, have been found to lack any form of NHEJ ^7^.

This study focuses on elucidating the phylogenetic distribution of the three distinct types of prokaryotic NHEJ pathways as outlined by Bertrand *et al*. ^8^ (Fig 1). These NHEJ classifications are based on the characterized pathways observed in *B. subtilis, Streptomyces ambofaciens*, and *Sinorhizobium meliloti*. In the case of *B. subtilis* it was determined that deletion of either the LigD or Ku protein hindered the functionality of NHEJ, suggesting both are necessary for the pathway ^5^. We will refer to this particular form of NHEJ that relies on only two proteins as “minimal NHEJ”. Further investigations conducted in *S. ambofaciens* revealed that mutations affecting other NHEJ-associated proteins, such as LigC, KuA, PolR, and PolK, had a significant impact on the overall efficiency of NHEJ ^9^. Here, the NHEJ pathway encompassing these four proteins will be designated as “core NHEJ”. More recent findings in *S. meliloti* unveiled the existence of two additional distinct functional NHEJ pathways, each activated under different stress conditions ^10^. The first pathway, operational in stress-free conditions, necessitates the presence of LigD2 and Ku2 proteins and will be referred to as the “main NHEJ” pathway. Conversely, the second pathway, induced exclusively under stress conditions, requires the participation of LigD4, Ku3, and Ku4 proteins and will be designated as the “secondary NHEJ” pathway. The combined occurrence of both main and secondary NHEJ pathways, as in *S. meliloti*, will be referred to as “multiple NHEJ” in this study.

**Fig 1.**
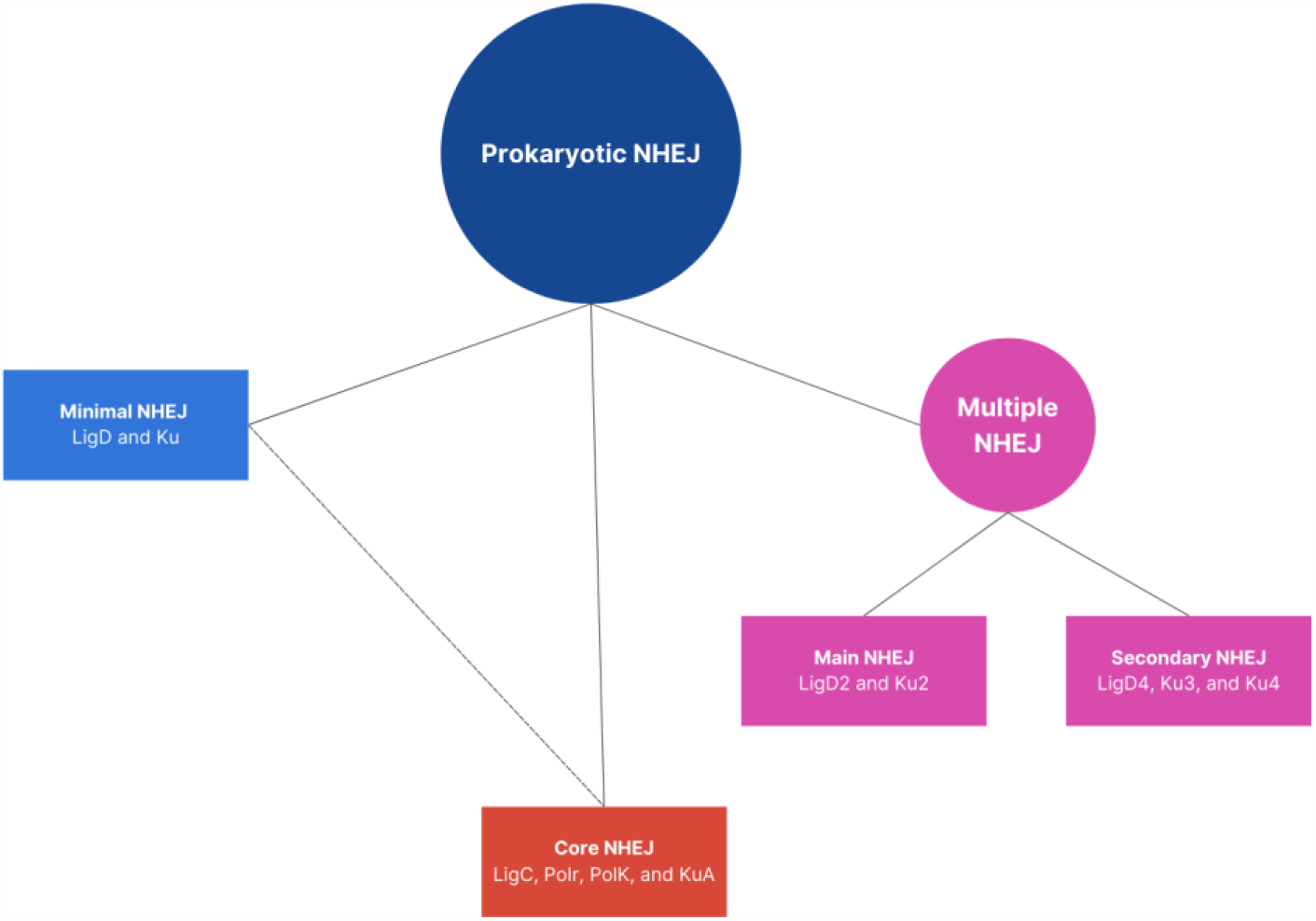
NHEJ pathways. A conceptual map illustrating the distinct categories of non-homologous end joining (NHEJ) observed in prokaryotes, accompanied by the corresponding protein constituents for each type. The prokaryotic NHEJ repertoire encompasses three major classifications: minimal, core, and multiple NHEJ. Minimal NHEJ requires LigD and Ku proteins. Core involves LigC, PolR, PolK, and KuA proteins. Multiple NHEJ comprises two additional pathways: main and secondary NHEJ. Main NHEJ requires LigD2 and Ku2 proteins, whereas secondary NHEJ relies on LigD4, Ku3, and Ku4 proteins.

Here, we perform a bioinformatic analysis on available bacterial and archaeal genome sequences to characterize the phylogenetic distribution of the various NHEJ pathways. We consider prokaryotic species with all proteins required for an NHEJ type to have a complete NHEJ pathway. To identify archaea species that may have prokaryotic NHEJ systems, we identified species with the four genes required for NHEJ as in *Methanocella pludicola* ^4^.

## METHODS

### Data

Computations were performed using the computer clusters and data storage resources of the HPCC, which were funded by grants from NSF (MRI-2215705, MRI-1429826) and NIH (1S10OD016290-01A1).

### Identification of Prokaryotes with a NHEJ Pathway

The UniProtKB database was used to BLAST the protein sequences of proteins necessary for functional prokaryotic NHEJ (LigD and Ku for minimal and core NHEJ, LigC, PolR, PolK, and KuA, for core NHEJ and LigD2, Ku2, LigD4, Ku3, and Ku4 for multiple NHEJ) using BLAST+ with the *E*-value cutoff of 0.0001 on the HPCC^8,11^. Query sequences were pulled from the NHEJ pathways of *Bacillus subtilis, Mycobacterium tuberculosis, Streptomyces ambofaciens*, and *Sinorhizobium meliloti* respectively. Results were downloaded and RegEx from Python was used to split the organism names from the results. Python was then used to remove duplicate organism names from the results and find duplicates in multiple results to sort organisms based on which NHEJ pathway they had.

### Identification of Archaea with a NHEJ Pathway

The UniProtKB database was again used to BLAST the sequences of proteins necessary for functional NHEJ using prokaryotic elements in *Methanocella paludicola* (Ku, Pol, PE, and Lig) using BLAST+ with the *E*-value cutoff of 0.0001 on the HPCC ^11,12^. Query sequences were pulled from *Methanocella paludicola*. Results were downloaded and RegEx from Python was used to split the organism names from the results. Python was then used to remove duplicate organism names from the results and find duplicates in multiple results to sort organisms based on if they had all the required protein sequences.

### Prokaryotic Phylogenetic Tree Construction

The 16S ribosomal ribonucleic acid (rRNA) sequences of identified prokaryotes were downloaded from the National Center for Biotechnology Information (NCBI)^13^. Single 16S rRNA sequences were chosen from prokaryotes with multiple sequences by selecting the sequence with the fewest Ns and longest sequence length. Multiple sequence alignments (msa) were generated using muscle v5 on default settings ^14^. IQTREE v2.2.0 was used to generate a maximum-likelihood (ML) phylogenetic tree ^15^. The best-fit model was found to be TIM3+F+R10 (LogL = -278,567.605, BIC = 582,595.148) using Model Finder ^16^. UFBoot2 (-bnni option) was used to assess branch supports ^17^. Finally, iTOL was used to visualize the trees and create colored ranges to signify NHEJ pathways ^18^.

### Archaeal Phylogenetic Tree Construction

The 16S ribosomal ribonucleic acid (rRNA) sequences of identified archaea were downloaded from the National Center for Biotechnology Information (NCBI)^13^. Single 16S rRNA sequences were chosen from archaea with multiple sequences by selecting the sequence with the fewest Ns and longest sequence length. Multiple sequence alignments (msa) were generated using muscle v5 on default settings ^14^. IQTREE v2.2.0 was used to generate a maximum-likelihood (ML) phylogenetic tree ^15^. The best-fit model was found to be GTR+F+R4 (LogL = -18,316, BIC = 38,109.029) using Model Finder ^16^. UFBoot2 (-bnni option) was used to assess branch supports ^17^. Finally, iTOL was used to visualize the trees ^18^.

## RESULTS

### Few Prokaryotes have Proteins Required for NHEJ

The protein sequences essential for NHEJ were subjected to Basic Local Alignment Search Tool (BLAST) analysis against the UniProtKB database ^11^. The obtained results were utilized to extract species names associated with individual proteins as well as those encompassing the entire complement of proteins necessary for each NHEJ type. Subsequently, the 16S ribosomal ribonucleic acid (rRNA) sequences of species possessing the complete set of NHEJ proteins were retrieved from the National Center for Biotechnology Information (NCBI) ^13^. These sequences were employed to generate an alignment and construct a phylogenetic tree.

Initially, this study focused on prokaryotes due to the existence of known (NHEJ) pathways as outlined by Bertrand *et al*. ^8^ (Fig 1). These pathways include minimal NHEJ, core NHEJ, and multiple NHEJ. Minimal NHEJ requires only two proteins: Ku and LigD. Core NHEJ requires more proteins: LigC, PolR, PolK, and KuA. Multiple NHEJ is further broken down into two subpathways: main NHEJ and secondary NHEJ. Main NHEJ is activated during times of stability and requires LigD2 and Ku2. Secondary NHEJ is activating during times of stress and requires LigD4, Ku3, and Ku4. Utilizing the information of these established pathways, an investigation was conducted to ascertain the number of prokaryotic species carrying all the essential genes necessary for at least one of the known functional NHEJ pathways.

It was found that prevalence of NHEJ in prokaryotes is relatively limited, with only 2624 out of 12,948 species (20.3%) available in the NCBI database exhibiting a complete form of NHEJ (Fig 2A) ^13^. Through this method it was found that NHEJ mechanisms are present in eight different prokaryotic classes (Fig 4). Unlike archaea, prokaryotic genomes tend to possess NHEJ protein coding sequences in close proximity as an operon, making it less common to encounter isolated components of the NHEJ machinery within prokaryotes ^4^.

**Fig 2.**
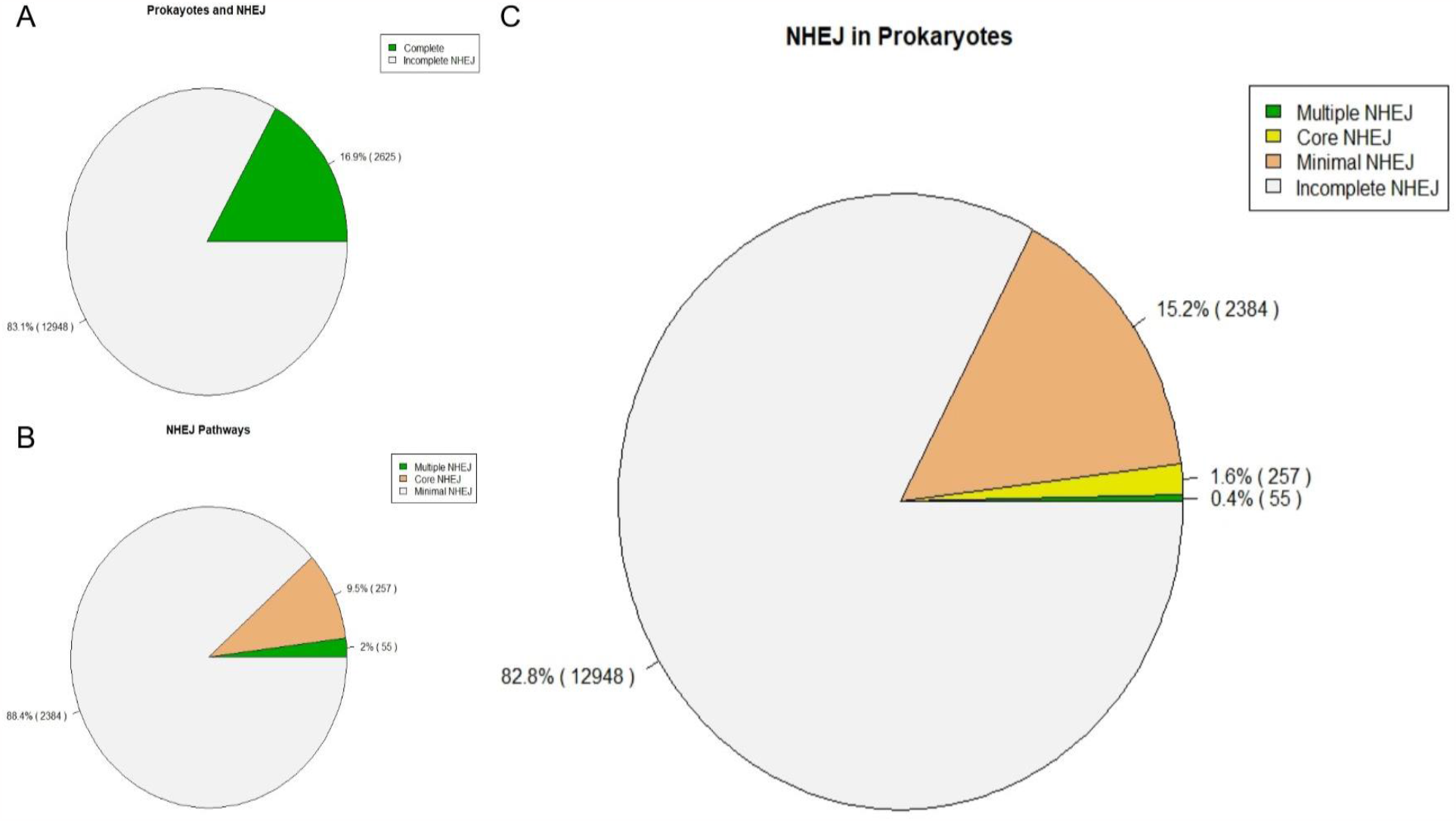
Prokaryotes and NHEJ. (**A**) The distribution of prokaryotic species possessing the complete set of proteins necessary for NHEJ is represented by green, whereas gray corresponds to species lacking one or more of the essential proteins. (**B**) Distribution of prokaryotes based on their NHEJ capabilities. The gray color represents prokaryotes lacking the NHEJ mechanism, while the orange color indicates prokaryotes with minimal NHEJ. Prokaryotes exhibiting core NHEJ are depicted by the yellow color, while the green color represents prokaryotes with multiple NHEJ. (**C**) Distribution of prokaryotes possessing distinct forms of NHEJ. The gray color represents the prokaryotes with minimal NHEJ, while the orange color indicates the prokaryotes with core NHEJ. The green color represents the prokaryotes with multiple NHEJ.

### Minimal NHEJ is the Most Commonly Found NHEJ Form in Prokaryotes

Following the determination of prokaryotic species with known NHEJ pathways, we further investigated the distribution of each established NHEJ pathway. Among the previously mentioned 2624 species possessing a complete set of genes required for NHEJ, an overwhelming majority of 2384 (18.4%) were found to be associated with the minimal NHEJ pathway, thus consolidating its prominence as the predominant form of NHEJ within the prokaryotic domain, as depicted in Figure 2C. A total of 20 prokaryotic species displaying minimal NHEJ were identified to also possess all the essential components necessary for multiple NHEJ, as illustrated in Figure 4.

Core NHEJ and multiple NHEJ genes are observed in only 2% of prokaryotic species with multiple NHEJ being the least common with it found in only 0.4% of prokaryotes (Fig 2B). Notably, it was observed that all prokaryotic organisms with 16S rRNA available on NCBI hosting the core NHEJ pathway genes also possess the multiple NHEJ pathway genes, indicating a consistent correlation between these two NHEJ pathways (Fig 4). Phylogenetic analysis showed the two pathways sporadically inherited, but notably, the genus *Streptomyces* was found to consistently carry core NHEJ. Core NHEJ and minimal NEHJ, on the other hand, were found to never occur together.

### Archaea Mostly Carry Parts of NHEJ

Since there is a prevalence of prokaryotic NHEJ components among archaeal species ^4^, a comparative analysis akin to the prokaryotic investigation was undertaken with archaea. The components of NHEJ found in *Methanocella paludicola* were considered required for a NHEJ pathway akin to that of prokaryotes to be carried out. The NHEJ proteins used in *M. paludicola* are Ku, Pol, PE, and Lig. The protein sequences of these proteins were used for a BLAST analysis against the UniProtKB database ^11^. Species names of carriers of all the proteins required for prokaryotic NHEJ were compiled. The 16S rRNA sequences of these were then retrieved from the NCBI database and utilized to create an alignment and construct a phylogenetic tree to showcase the evolutionary relationship of archaea carrying prokaryotic NHEJ ^13^.

Among the 567 Archaea species documented in the NCBI database, only 16 do not have any prokaryotic NHEJ genes (Fig 3A). However, of the remaining 545 species, only a total of 97 species from six different classes possess all the requisite parts for NHEJ indicating that while many Archaea carry parts of NHEJ genes, few could carry out the prokaryotic NHEJ pathway (Fig 3B). The phylogenetic relationships of the 16S rRNA from species with all the requisite parts for NHEJ is shown in Figure 5. In the case of Asgard lineages, while they do harbor certain constituents of the NHEJ system, they lack the entirety of the required elements for functional NHEJ.

**Fig 3.**
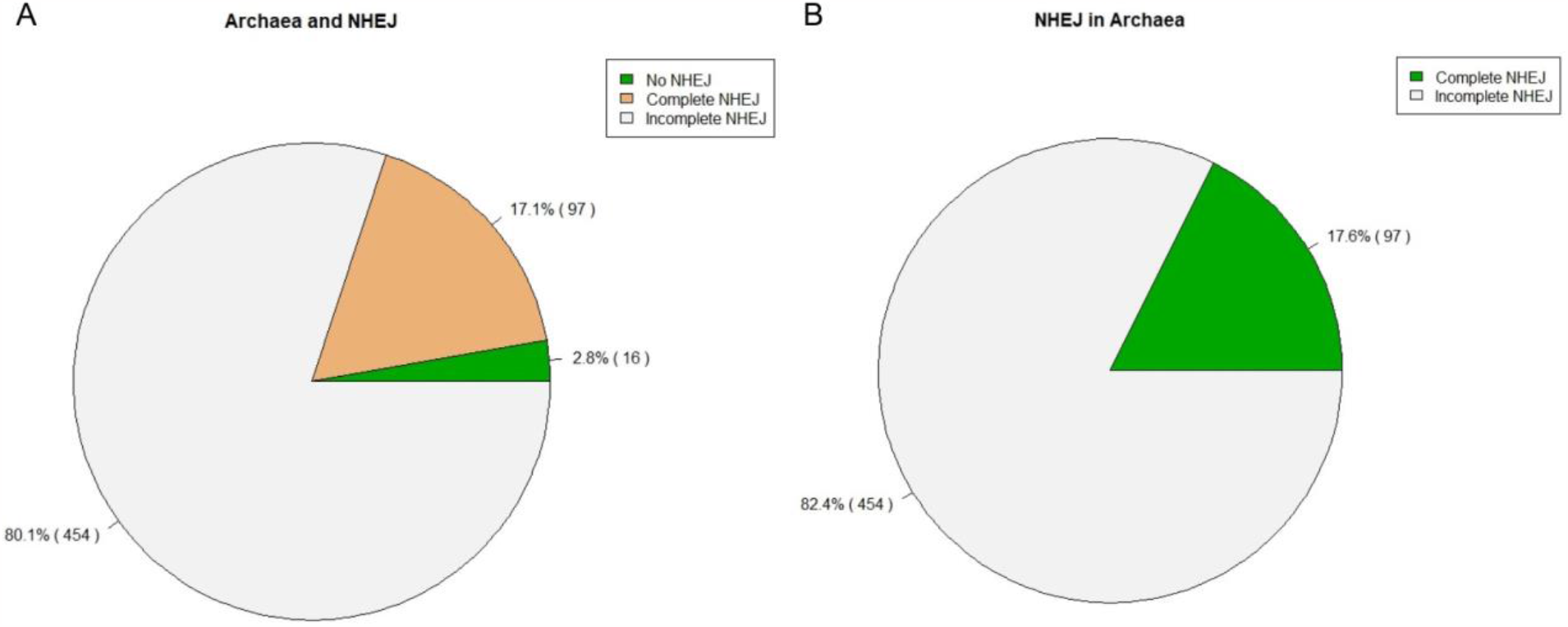
Archaea and NHEJ. (**A**) The distribution of archaeal species exhibiting various levels of prokaryotic NHEJ. The gray shade represents the number of archaeal species possessing at least one prokaryotic NHEJ protein. The orange shade corresponds to the number of archaeal species encompassing all the proteins essential for prokaryotic NHEJ. Finally, the green shade represents the number of archaeal species lacking any prokaryotic NHEJ proteins. (**B**) Distribution of archaeal species with respect to carrying prokaryotic NHEJ. The gray shade corresponds to the number of archaeal species harboring at least one prokaryotic NHEJ protein. Conversely, the green shade represents the number of archaeal species possessing all the requisite prokaryotic NHEJ proteins necessary for executing the pathway.

**Fig 4.**
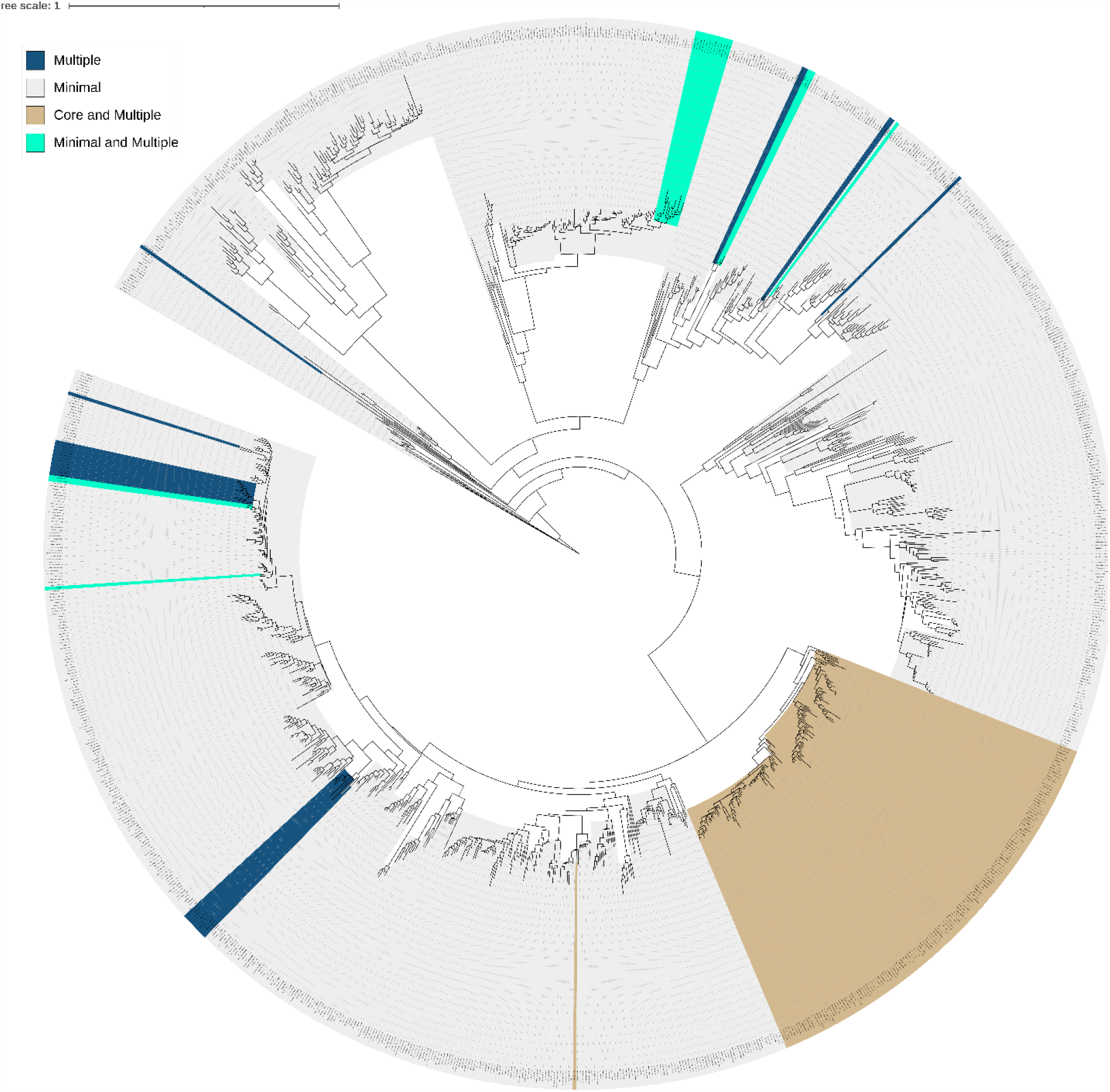
NHEJ Pathways in Prokaryotes. A phylogenetic tree of prokaryotic species exhibiting distinct forms of NHEJ. The tree scale bar signifies the percent difference between the 16S rRNA sequences. The brown color signifies prokaryotes that possess both core and multiple NHEJ pathways, while the dark blue color designates prokaryotes exclusively harboring the multiple NHEJ pathway. Additionally, the gray color highlights prokaryotes exclusively characterized by the presence of the minimal NHEJ pathway. Finally, the cyan signifies the prokaryotes that carry the genes for both the minimal NHEJ and multiple NHEJ pathways.

**Fig 5.**
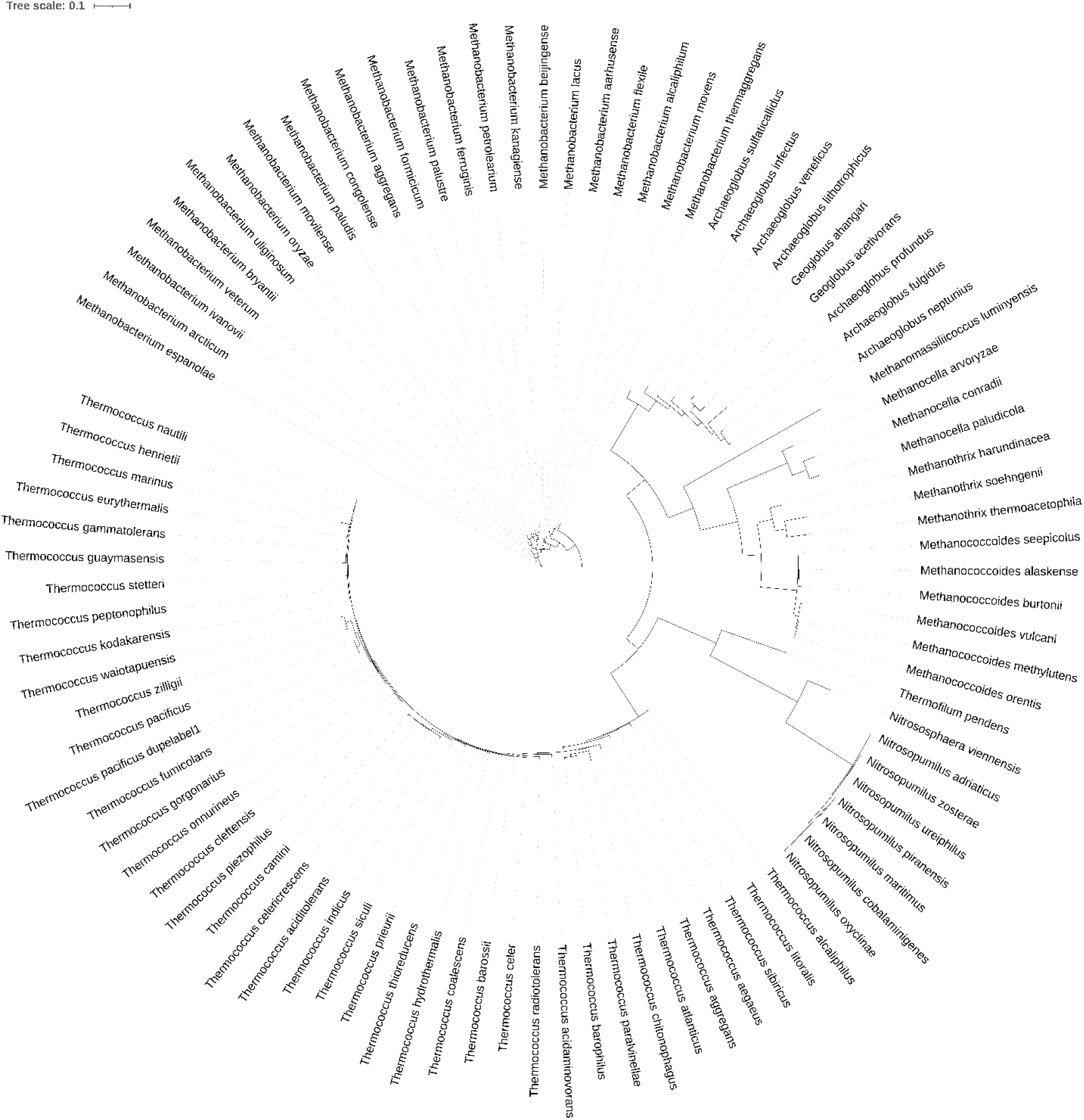
NHEJ in Archaea. Phylogenetic tree that illustrates the evolutionary relationships among archaeal lineages with NHEJ. The tree scale bar signifies the percent difference between the 16S rRNA sequences.

## DISCUSSION

To identify prokaryotic species with a higher likelihood of possessing a functional NHEJ pathway, a collection of prokaryotic species encompassing all the requisite proteins for NHEJ pathways was compiled. The NHEJ pathways that were used were minimal NHEJ, core NHEJ, and multiple NHEJ, as described in Bertrand et al. 2019^8^. For minimal NHEJ, the essential proteins were LigD and Ku, whereas LigC, KuA, PolR, and PolK were required for core NHEJ. Multiple NHEJ comprises two subpathways, namely main NHEJ and secondary NHEJ, necessitating LigD2 and Ku2 for the former and LigD4, Ku3, and Ku4 for the latter. Species possessing all the proteins required for a particular pathway were classified as having “complete NHEJ” and thus potentially capable of executing the pathway. For archaea, species with PE, Lig, Pol, and Ku were considered ‘complete’ as those are the only NHEJ proteins that have been found in archaea thus far ^4^.

Our study revealed that while 80% of prokaryotes do not have all the proteins required for functioning NHEJ, minimal NHEJ is the most common form found (Fig 2A). Notably, minimal NHEJ only relies on two distinct proteins, whereas the other prokaryotic NHEJ forms involve a greater number of proteins, suggesting that the simplicity of minimal NHEJ may facilitate its widespread occurrence.

A limited number of species exhibiting minimal NHEJ also possess the requisite components for multiple NHEJ, as depicted in Figure 4. The observed distribution appears to be attributed to horizontal gene transfer (HGT) rather than vertical gene transfer, considering the considerable phylogenetic distance separating the species on the phylogenetic tree.

The least commonly observed NHEJ form in prokaryotes is multiple NHEJ, which comprises main and secondary NHEJ subpathways. Intriguingly, species exhibiting multiple NHEJ could also possess core NHEJ (Fig. 4). Core NHEJ was also exclusively identified in the *Streptomyces* genus, implying a potential vertical transfer of core NHEJ in that genus. Furthermore, the occurrence of both multiple and core NHEJ in similar species across different clades suggests a combination of horizontal and vertical gene transfers for these NHEJ forms. Reliance of core NHEJ on multiple NHEJ could also be implied by this. It is possible that core NHEJ is also an accessory part of multiple NHEJ or that the two share similar mechanisms and thus genetics as well. Since core and minimal NHEJ were found to never occur together, it is possible that one inhibits the other leading to the organism choosing one pathway over the other.

Based on our results, it is plausible that the majority of archaea actually carry at least one protein involved in prokaryotic NHEJ (Fig 3A). However, only approximately 18% of archaea harboring an NHEJ protein possess all the proteins necessary for functional prokaryotic NHEJ. Although these archaea possess all the requisite proteins, their inability to execute the pathway may be attributed to the lack of operon fusion, as observed in prokaryotes ^4^. This could potentially be attributed to horizontal gene transfer events involving individual protein sequences from prokaryotes.

Interestingly, while Asgard lineages exhibit partial prokaryotic NHEJ components, none were found to possess all the required protein sequences. This observation suggests that horizontal gene transfer between prokaryotes or archaea and Asgard lineages may not be as straightforward. Alternatively, it raises the possibility that asgard lineages may rely upon eukaryote or eukaryote-like NHEJ systems rather than prokaryotic.

## Notes

### Competing Interest Statement

The authors have declared no competing interest.

